# Computer assisted multi-level optimization of malonyl-CoA availability in *Pseudomonas putida*

**DOI:** 10.1101/2024.11.20.624107

**Authors:** Christos Batianis, Rik P. van Rosmalen, Pedro Moñino Fernandez, Enrique Asin-Garcia, Maria Martin-Pascual, Markus Jeschek, Ruud Weusthuis, Maria Suarez-Diez, Vitor A. P. Martins dos Santos

**Affiliations:** Laboratory of Systems and Synthetic Biology, Wageningen University & Research, Wageningen, 6708 WE, The Netherlands; Bioprocess Engineering, Wageningen University & Research, Wageningen, 6708 PB, The Netherlands; Department of Biosystems Science and Engineering, ETH Zurich, CH-4058 Basel, Switzerland; Institute of Microbiology, Synthetic Microbiology Group, University of Regensburg, Regensburg D-93053, Germany; LifeGlimmer GmbH, Berlin, 12163, Germany

**Keywords:** *Pseudomonas putida*, Malonyl-CoA, Genome-scale modeling, CRISPRi, RBS optimization

## Abstract

Malonyl-CoA is the major precursor for the biosynthesis of diverse industrially valuable products such as fatty acids/alcohols, flavonoids, and polyketides. However, its intracellular availability is limited in most microbial hosts, hampering the biological synthesis of such chemicals. To address this limitation, we present a multi-level optimization workflow using modern metabolic engineering technologies to systematically increase the malonyl-CoA levels in Pseudomonas putida. The workflow involves the identification of gene downregulations, chassis selection, and optimization of the acetyl-CoA carboxylase complex through ribosome binding site engineering. Computational tools and high-throughput screening with a malonyl-CoA biosensor enabled the rapid evaluation of numerous genetic targets. Combining the most beneficial targets led to a 5.8-fold enhancement in the production titer of the valuable polyketide phloroglucinol. This study demonstrates the effective integration of computational and genetic technologies for engineering P. putida, opening new avenues for the development of industrially relevant strains and the investigation of fundamental biological questions.

## 1. INTRODUCTION

Biobased production of bulk and fine chemicals is essential in our quest to generate sustainable alternative to petrochemical-based processes. Currently, biobased products are utilized across a range of industries, including pharmaceuticals, food, and agriculture (Dahiya et al., 2020). Metabolic engineering and synthetic biology have played a pivotal role in the optimization of produc-tion lines and successful commercializion of biobased chemicals (Voigt, 2020; Liu et al., 2023). Advances in technologies such as DNA synthesis/sequencing, metabolic modeling, artificial intelligence and robotics, enable faster and more efficient development of industrially attractive strains (Ko et al., 2020). A crucial aspect of metabolic engineering involves tailoring microbes’ central carbon metabolism to increase availability of key precursor molecules that serve as universal building blocks for numerous relevant bioconversions. One such important precursor is malonyl-CoA, which plays a central role in several biologically essential processes, including fatty acid biosynthesis. Malonyl-CoA also serves as a building block for the production of diverse valuable compounds such as polyketides, flavonoids, stilbenes, and coumarins (Milke & Marienhagen, 2020). Often, these compounds are secondary metabolites in plants or microbes; however, due to the inherent disadvantages associated with their natural producers, increasing attention has been paid to the development of heterologous microbial cell factories towards their economically viable production (Johnson et al, 2017).

While malonyl-CoA-derived products have been successfully produced using various microbial chassis, the endogenous levels of malonyl-CoA are typically tightly regulated and maintained at low concentrations, hindering production efficiency and commercial viability (Milke & Marienhagen, 2020). Hence, increasing its intracellular pool is a common challenge in metabolic engineering. Current approaches to tackle this challenge focus on rational engineering, dynamic (autonomous or non-autonomous) regulation of the malonyl-CoA levels and/or the exploitation of various biosensors (Johnson et al, 2017; Dinh & Prather, 2019; Zhou et al., 2021). Genetic engineering tools, such as CRISPR, CRISPRi/a, and sRNAs have been instrumental in generating mutant libraries for high-throughput screening of promising genetic targets, accelerating the Design-Build-Test-Learn cycle framework (Liu et al., 2018; Tarasava et al., 2018; Ferreira et al., 2019). However, given that thousands of genetic elements could serve as potential targets, it remains challenging to identify which targets need to be regulated and how. While genome-scale target identification is possible through recent advances in robotic automation, it can be prohibitively expensive and unavailable in many cases. As such, implementing computational tools into the workflow offers an excellent alternative that helps narrow down candidate targets (Xu et al., 2011; Tan et al., 2018, Moreno-Paz et al., 2024). In this context, constraint-based models have been widely employed as a powerful tool for understanding and re-design of microbial metabolism, allowing for the prediction of metabolic scenarios favoring the production of specific metabolites (Bordbar et al., 2014; Machado & Herrgård, 2015). These scenarios can be rapidly evaluated with high-throughput genome engineering tools and biosensors, enabling medium-to high-throughput screening experiments, while successful strains are further improved during subsequent rounds of computation and optimization.

Most of the available malonyl-CoA biosensors and genetic tools have been optimized for a small number of generic microbial chassis, mainly confined to *Escherichia coli* and *Saccharomyces cerevisiae* (Johnson et al., 2017). However, several promising microbial chassis, such as *Corynebacterium glutamicum*, *Streptomyces coelicolor*, *Yarrowia lipolytica*, and *Pseudomonas putida*, exhibit unique features to produce malonyl-CoA products (Milke et al., 2019; Ma et al., 2020; Mezzina et al., 2021; Li et al., 2021). *P. putida* KT2440 is a Gram-negative soil bacterium, HV1 certified (Kampers et al., 2019), with numerous characteristics that render it an attractive host for industrial biocatalysis (Martins Dos Santos et al., 2004; Poblete-Castro et al., 2012). *P. putida* is also endowed with several traits required for the production of malonyl-CoA-derived products. For instance, it has been extensively employed as a microbial cell factory of oleochemicals such as α,ω-diols (Lu et al., 2021) and polyhydroxyalkanoates due to its natural ability to accumulate medium-chain-length polyhydroxyalkanoates (mcl-PHAs) as carbon storage under nutrient limitation (Mezzina et al., 2021). In addition to oleochemicals, it is also an attractive host for producing natural products such as polyketides and non-ribosomal peptides (Loeschcke & Thies, 2015). *P. putida* is equipped with a wide substrate range phosphopantetheinyl transferase (PPTase) that is often more suitable than *E. coli’s* PPTases, bypassing the need of additional introduction of foreign PPTase genes (Loeschcke & Thies, 2015). Additionally, the availability of an extensive genetic toolbox and genome-scale metabolic models reinforce potential for implementation of *P. putida* as a platform for industrial biotransformations (Martin Pascual et al., 2021). Nevertheless, to unlock the potential of *P. putida* as an effective producer of malonyl-CoA-derived products, it is essential to increase its intracellular malonyl-CoA levels, which, in turn, have been shown to be relatively low compared to other hosts (Gläser et al., 2020).

In this work, we present a multi-level optimization workflow using a series of modern metabolic engineering technologies to systematically engineer the intracellular malonyl-CoA levels in *P. putida* (Fig. 1). Our approach involves the concurrent execution of three optimization levels: identification of gene knockdown targets, chassis selection, and ribosome binding site (RBS) optimization of the acetyl-CoA carboxylase complex. Computational methods guided the selection of potential engineering options, which were subsequently evaluated using an extracellular malonyl-CoA biosensor (Fig. 1). All predicted genetic interventions were rapidly applied using CRISPRi and Cre-mediated high-throughput gene integration. Through this approach, we identified gene knockdown targets and an optimal ACC RBS combination that significantly increased the biosensor signal up to 5.4-and 2-fold, respectively. Finally, the most beneficial genetic targets resulted from the screening were applied in combination to significantly increase the production titer of the valuable polyketide phloroglucinol by 5.8-fold. This tailored *P. putida* strain could be highly useful for enhanced bioproduction of any valuable product where the cellular malonyl-CoA abundance is the limiting step. Moreover, the current study provides technical insights into the application of novel computational and genetic technologies in *P. putida* and paves the way for their functional implementation in the development of industrially relevant strains, as well as for the investigation of fundamental biological questions.

**Figure 1.**
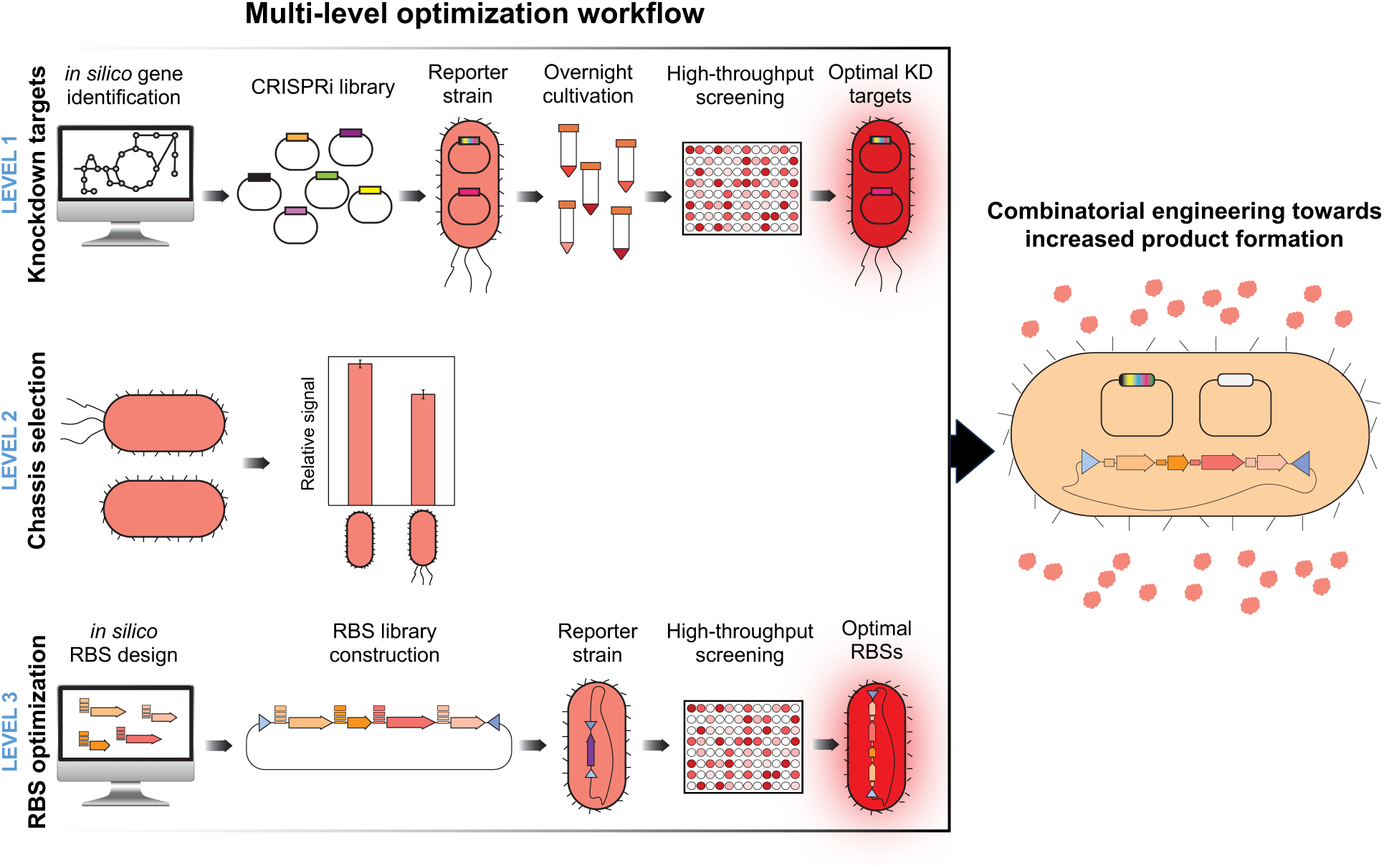
Multi-level optimization workflow to increase the intracellular malonyl-CoA levels in *P. putida*. LEVEL 1: Gene knockdown targets were predicted *in silico* using a newly developed flux sampling pipeline. A CRISPRi library comprised of all selected targets was transformed in the reporter strain carrying the malonyl-CoA biosensor and promising gene knockdowns were identified based on the relative signal using a microplate reader. LEVEL 2: The wild-type strain *P. putida* KT2440 was compared with the streamlined variant *P. putida* EM42 using the malonyl-CoA biosensor for their ability to produce malonyl-CoA. LEVEL 3: A combinatorial ACC RBS library was designed *in silico* and constructed in a synthetic *accABDC* operon. The operon library was integrated directly into the chromosome of the reporter strain and the optimal RBS combination was identified based on the relative signal. The most beneficial genetic interventions were applied in combination to enhance the production of the malonyl-CoA-derived product phloroglucinol.

## 2. MATERIAL AND METHODS

### 2.1 Bacterial strains and plasmids

A list of the bacterial strains and plasmids used in this study is presented in Table 1.

**Table 1.**
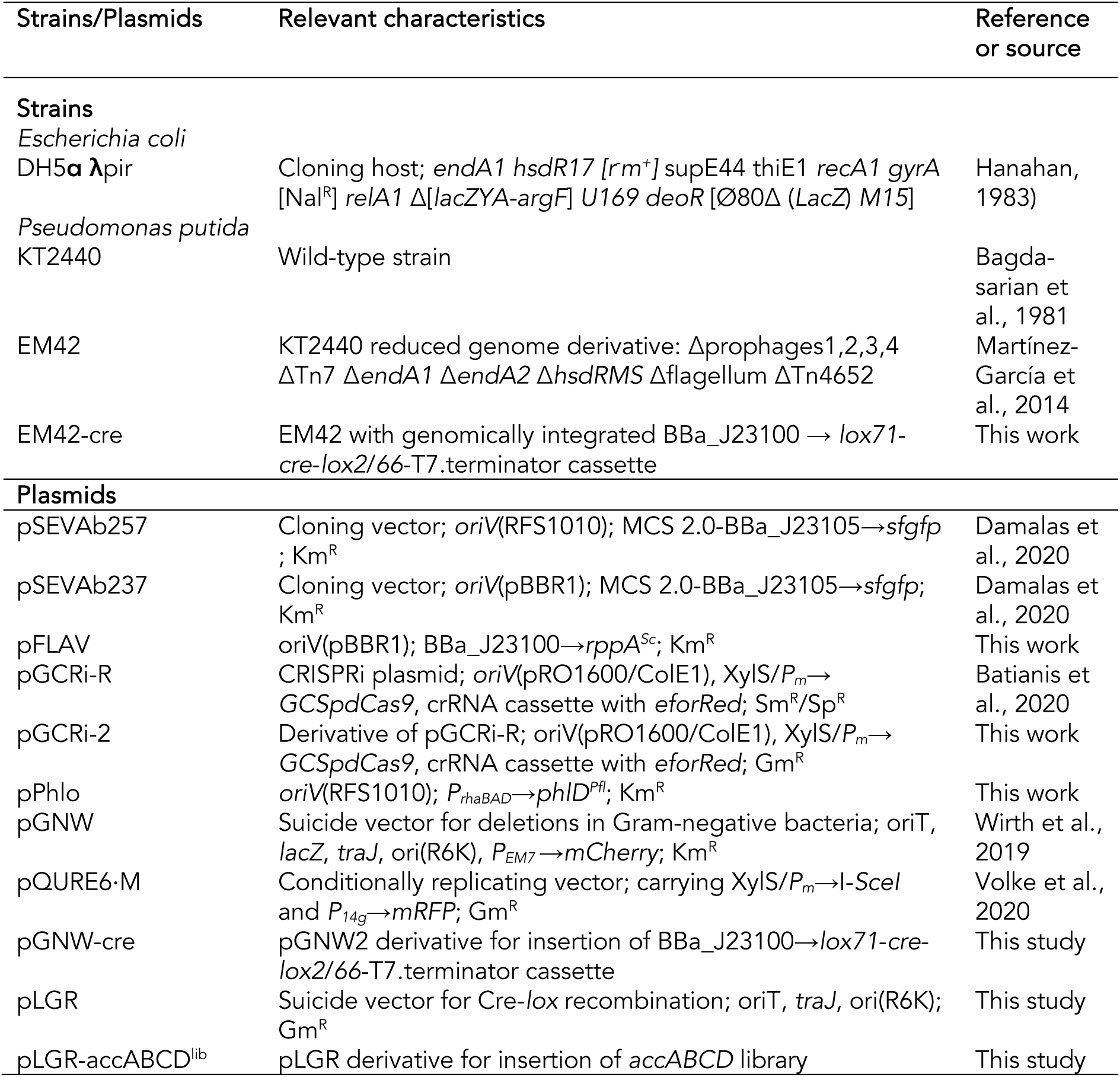
Strains and plasmids used in this study.

### 2.2 Culture medium and conditions

*E. coli* and *P. putida* strains were grown in Lysogeny Broth (LB) medium containing 10 g/l tryptone, 10 g/l NaCl and 5 g/l yeast extract, unless otherwise indicated and incubated at 30 °C and 37 °C, respectively. In case of solid media 1.5% (w/v) agar was added. Antibiotics were supplemented to all media whenever required at concentrations of 50 μg/ml for kanamycin (Km) and 10 μg/ml for gentamycin (Gm). All falcon tube experiments for flaviolin production were performed in MOPs minimal medium comprised of 0.29 mM K_2_SO_4_, 32.5 μM CaCl_2,_ 1.32 mM K_2_HPO_4_, 8 μM FeCl_2_, 40 mM MOPS, 4 mM tricine, 9.52 mM NH_4_Cl, 0.01 mM FeSO_4,_ 0.52 mM MgCl_2_, 0.03 μM (NH_4_)_6_Mo_7_O_24_, 4μM H_3_BO_3_, 50 mM NaCl, 0.3μM CoCl_2_, 0.1 μM CuSO_4_, 0.8 μM MnCl_2_, and 0.1 μM ZnSO_4_ (LaBauve & Wargo, 2012). Plasmids pGCRi-2 and pPhlo were induced using 1 mM of 3-methylbenzoate (3MB) and 4 mM of L-rhamnose, respectively. Flask cultures were performed in 250 ml Erlenmeyer flasks filled with 25 ml of MOPs minimal medium supplemented with 70 mM of glucose and incubated with rotary shaking at 200 rpm. Flasks were inoculated to OD_600_ = 0.2 from an overnight LB culture and product concentrations were determined at the end of the experiment.

### 2.3 Predicting knockdown targets through metabolic flux sampling

The iJN1462 genome-scale metabolic reconstruction (Nogales et al., 2019) was used for all simulations, using glucose as a carbon source at a maximum uptake rate of 6.3 mmol/gDW/h and unlimited uptake of all other essential nutrients. All reactions were constrained to a maximum absolute flux of 1000 mmol/gDW/h. Simulations were performed using the OptGPSampler (Megchelenbrink et al., 2014) implementation in the COBRApy (Ebrahim et al., 2013) library for flux balance analysis (version 0.16.0), utilizing the Gurobi optimizer (version 8.1.1). For each condition, fluxes were sampled 10^6^ times with a thinning factor of 100, leading to a final number of 10^4^ samples per condition. Convergence was verified using the Geweke diagnostic (Geweke, 1991). Additional genome annotation was retrieved from the Pseudomonas Genome Database (pseudomonas.com). The source code of the complete analysis is available as open-source information at https://gitlab.com/wurssb/pputida-mcoa-optimization. To compare the flux distributions under growth and production conditions, flux distributions sampled in three scenarios were compared. First, the growth condition was simulated, constrained by a minimum biomass flux of 90% of the optimal biomass flux. In order to increase the reliability of the flux profile, a parsimonious FBA (pFBA) (Lewis et al., 2010) constraint was added, under the assumption that enzyme usage under optimal growth conditions is well-optimized through evolution. This was implemented by constraining the maximum of the sum of all absolute fluxes to be less than 125% of the minimum sum of the absolute fluxes at the optimal growth rate. Second, the production condition was simulated by constraining product formation to be at least 90% of the maximum possible production.

After sampling each of the three scenarios, several criteria were applied to filter out the most promising reactions to knockdown. Reactions were kept only if they met all the following criteria.

- The reaction has genes annotated but is not annotated with multiple isozymes.
- The significance value of a 2-sample Kolmogorov-Smirnov test between the production and the growth cases, and the production and the slow growth cases, was less than 0.05 after multiple testing correction using the Bonferroni method. However, due to the large sample sizes the Kolmogorov-Smirnov test is overpowered, and therefore not very distinctive. Thus, the KS-statistic was additionally required to be more than 0.5 between the growth and production case and 0.25 between the slow growth and production case.
- The absolute difference between the mean of the production and the growth flux profiles is more than 10^-2^ mmol/g dry weight/h.
- The reaction flux had a correlation with the biomass reaction flux lower than 0.9 when considering all the samples in all possible scenarios.

As a final selection criterium, reactions were clustered based on the correlation of the samples of the production phenotypes, using average based clustering on the Euclidean distance of the correlation profiles. This helped to group reactions in related pathways or processes together in order to select targets in different pathways and explore a more diverse set of possible interventions. Based on the largest fold change between the producing and growth state, potential target reactions were then selected and further curated by expert knowledge. Finally, potential knockdown targets for these reactions were verified not to be involved in other reactions requiring over-expression in the production scenario.

For the selection of dual mutants, all reactions having at least one of their required genes as a target for this first knockdown were limited to an absolute flux less than the 25^th^ percentile of the wild-type flux distribution during growth. Analyses were then performed similarly to the wild-type case described above. In addition, an additional analysis was performed where the effect of the second knockdown on the reaction flux was compared to the effect of the same knockdown on the wild-type strain. If the effect of the second knockdown (i.e., the difference between the production and the growth flux profile) was greater in the strain with both knockdowns applied than in the wild-type strain with only the second knockdown applied, the pair of reactions was classified as synergistic. Conversely, if the effect of the second knockdown was larger in the wild-type strain than in the first knockdown strain, the reaction pair was classified as anti-synergistic.

### 2.4 RBS prediction and smart library design

To optimize absolute and relative expression levels of the synthetic *accABCD* operon, we constructed a combinatorial RBS library based on *in silico* predictions and library design. First, we used a previously developed deep learning model to predict the strength of 65536 RBS variants for each of the four genes (Höllerer et al., 2020). More precisely, we allowed full randomization of eight consecutive nucleotides in the 5’-UTR in a distance of 5 base pairs to the respective start codon (*accA*: AATTNNNNNNNNAATTA, *accB*: ATATNNNNNNNNATATA, *accC*: TAATNNNNNNNNATATT, *accD*: TTAANNNNNNNNTTAAT). The resulting prediction data were then reduced using the RedLibs tool (Jeschek et al., 2016) to obtain libraries of six RBSs encoded by a single degenerate sequence for each gene (*accA*: AATTGAGVGAGKAATTA, *accB*: ATATAVGGARACATATA, *accC*: TAATTSGGGBGTATATT, *accD*: TTAACAAGRGBGTTAAT). These libraries uniformly span the accessible range of RBS strengths enabling effective coverage of the multidimensional expression level space whilst keeping the screening effort manageable (theoretical combinatorial size: 6^4^ = 1296 variants).

### 2.5 General cloning procedures

All primers used in this study are listed in the Supplementary Materials. DNA fragments for plasmid construction were amplified with Q5 High-Fidelity DNA polymerase (New England Biolabs) following the procedures described in the SevaBrick assembly protocol (Damalas et al., 2020). The fragments were separated by electrophoresis in 1% (w/v) agarose gel and PCR purification was performed with the NucleoSpin PCR purification kit (Macherey-Nagel) according to the manufacturer’s instructions. DNA assembly was carried out by SevaBrick assembly using *Bsa*I restriction enzyme and T4 ligase (New England Biolabs) as described by Damalas et al., 2020. Transformations were performed in *E. coli* DH5α chemically competent cells as described by Green and Rogers, 2013. For transforming *P. putida*, electrocompetent cells were prepared from an overnight culture by washing the cell pellet three times with 1 ml of 300 mM sucrose, according to Choi et al., 2006. Electroporation was carried out in a Gene Pulser/Pulse Controller (Bio-Rad Corp.) with a voltage of 2.5 kV, 25 μF capacitance and 200 Ω resistance. After transformation, cells were plated on selective LB-agar plates with the appropriate antibiotics and single colonies were subjected to colony PCR with Phire Hot Start II DNA polymerase (Thermo Fisher Scientific). The correct colonies were grown overnight in 50 ml falcon tubes with 10 ml of LB medium supplemented with the corresponding antibiotics. The plasmids were extracted from the overnight cultures with the GeneJET Plasmid Miniprep Kit (Thermo Fisher Scientific) and sent for sequencing (MACROGEN Inc.). For the construction of pFLAV, the codon-optimized *Streptomyces coelicolor* A3(2) *rppA* was ordered from IDT (Integrated DNA Technologies), domesticated into the repository vector pSB1C3, and subsequently cloned into a pre-linearized pSEVAb23 backbone under the control of the BBa_J23100 promoter by SevaBrick assembly (Damalas et al., 2020). The same procedure was followed for the construction of pPhlo using the codon-optimized *Pseudomonas fluorescens* Pf-5 *phlD* gene and the pSEVAb25 backbone. Marker-less gene integration was facilitated by homologous recombination using the pGNW suicide vector as described by Volke et al., 2020 and Wirth et al., 2019.

### 2.6 CRISPRi library construction

For the CRISPRi-mediated screening, we utilized the pGCRi plasmid with minor modifications (Batianis et al., 2020). The new pGCRi-2 plasmid carries the gentamycin instead of the streptomycin resistance gene as it was observed that *P. putida* frequently gains resistance over streptomycin making the selection process complicated. The assembly of the target spacers was performed using Golden Gate following the procedure described by Batianis et al., 2020. All spacers were designed to target the promoter sequence or the ATG region of each *in silico* predicted gene target. The latest annotated genome of *P. putida* KT2440 was used as the reference sequence for promoter prediction using the online tool Softberry BPROM (Solovyev & Salamov, 2011). When the promoter could not be predicted, or the genes were inside an operon, the ATG region was targeted instead. As a control, a non-targeting spacer for both *P. putida* and *E. coli* was designed. The full list of spacers is displayed in the Supplementary Materials. All spacer oligonucleotides were purchased from IDT (Integrated DNA Technologies).

### 2.7 Construction of the *accABCD* RBS library

Following the *in silico* library design, each gene was PCR amplified with degenerate forward oligonucleotides encoding all predicted RBS combinations. Both forward and reverse oligonucleotides contained *Bsa*I restrictions sites. In the degenerate forward primer, two nucleotides within the RBS sequence were changed to all possible combinations. To increase the efficiency of the final assembly, we performed two rounds of ligation. First, the amplification products of *accA* and *accB* were ligated together linearly, as well as those of *accC* and *accD*. Next, we reamplified the linear ligation products and assembled them together into the pLGR backbone using Golden Gate.

### 2.8 Screening of CRISPRi library using the RppA-sensor

Colonies of *P. putida* mutants carrying pFLAV were picked and cultured overnight in 50 ml tubes with vent caps containing 10 ml of LB medium and the required antibiotics. The overnight cultures were centrifuged for 10 minutes at 4700 rpm, the supernatant was discarded, and the cells were resuspended in MOPS medium. The resuspended cells were diluted to OD_600_ = 0.2 into 14 ml falcon tubes with MOPS medium containing 70 mM of glucose and antibiotics. The fresh cultures were incubated overnight at 30 °C and 200 rpm rotary shaking. After 24 h, 1 ml of each culture was transferred into a 1.5 ml eppendorf tube, and cells were harvested by centrifugation at >11000 g for 2 min. Next, the supernatant containing flaviolin was transferred into another 1.5 ml eppendorf tube and cells were resuspended in 1 ml MOPS. The flaviolin in the supernatant and the cell density were measured in triplicate at OD_340_ and OD_600_, respectively, in 96-well plates using the Synergy microplate reader (Biotek Instruments). The OD_600_ of the resuspended cells was determined after 4-fold dilution in MOPS medium.

### 2.9 Screening of RBS libraries using the RppA-sensor

*P. putida* EM42 carrying pFLAV and the genomically integrated *lox*-*cre*-*lox* cassette was transformed with the pLGR-accABCD library and plated in LB plates containing gentamycin and kanamycin. After plating, 2000 colonies were picked and inoculated in deep 96-well plates containing 200 μl of LB medium and the required antibiotics. The plates were incubated at 30 °C in rotary shaking 200 rpm. After 24 h, cells were pelleted at 4500 rpm for 10 min, the supernatant was transferred into 96-well plates and the cell pellet was resuspended in PBS buffer and later transferred into 96-well plates. The relative flaviolin production was determined as previously described.

### 2.10 Analytical methods

The concentration of phloroglucinol was analyzed by high-performance liquid chromatography (HPLC). Briefly, samples were centrifuged at 12000 g for 5 min, and then the supernatants were filtered through a 0.20 μm nitrocellulose filter and analyzed by an HPLC (Thermo Fischer Scientific) equipped with UV/vis. The column (C18, 5 µm, 250 × 4.6 mm, Agilent, USA) was eluted at 30 °C using acetonitrile/water (4/6, v/v) as the mobile phase at a flow rate of 1 ml/min. The detection wavelength was 254 nm. A phloroglucinol calibration curve was prepared using pure standards (99% purity) purchased from Sigma-Aldrich.

## 3. RESULTS

### 3.1 LEVEL 1: Identification of gene knockdowns

#### 3.1.1 In silico selection of putative gene knockdown targets

To predict putative knockdown targets for increased malonyl-CoA availability, the *P. putida* metabolic flux state during growth was compared to the metabolic state of *P. putida* when overproducing malonyl-CoA. Reactions were identified as potential overexpression (Fig. 2B) or downregulation (Fig. 2C) targets using the difference between the *in silico* flux distributions during growth and production. If the flux of a reaction is significantly different between the two conditions, it indicates that modulating the flux through knockdown or overexpression could be required to shift from the growth to the production phenotype. Since the number of reactions with a significant difference between growth and production was high, several criteria were used to filter down the number of reactions (Fig. 2G). The first criteria included the presence of annotated genes in the model and the absence of iso-enzymes. Next, the targets were screened for a significant difference in flux between the growth and production conditions. However, reactions can differ in flux not only because they are required to be high or low in one of the scenarios but also because they are less constrained in either scenario. Thus, an additional reference scenario, low growth, was simulated to discern between these fluxes. This scenario was implemented by constraining the maximum growth rate to be the same as the maximum growth rate possible in the production scenario. Furthermore, reactions with a flux correlating strongly to the biomass flux were eliminated to avoid targets more likely to be related to the low growth rates observed in the production phenotype than the high production rate. After applying these criteria as well as eliminating reactions with a small absolute difference in flux between the conditions, 250 reactions remained, belonging to 147 correlated groups. Genes catalyzing these reactions (221) were manually curated as potential targets, for example, by screening the genes for potential off-target effects on other reactions than the intended target. In addition to the *in silico* predicted targets, several genes identified through rational criteria and literature review were added to the list, such as targets to increase ATP availability. Several genes in the TCA cycle were added even though they did not match all criteria, as we hypothesized that they could increase acetyl-CoA availability which in turn could lead to an increase in the malonyl-CoA pool. The final selected subset, along with the selection criteria, is presented in the Supplementary Materials. Next, pairwise combinations of a selection of successful targets were identified by comparing the difference in fluxes between growth and production in the wild-type strain versus the simulated mutant strain (Fig. 2D-F), where the mutant was simulated by constraining the flux through affected reactions to be less than the 25_th_ percentile of the flux sample from the wild-type strain. Several synergistic targets, as well as new targets not included in the single target analysis were found, although testing the target combinations experimentally was not in the scope of the current work. The complete dataset of both single and pairwise targets can be found in the repository at https://gitlab.com/wurssb/pputida-mcoa-optimization.

**Figure 2.**
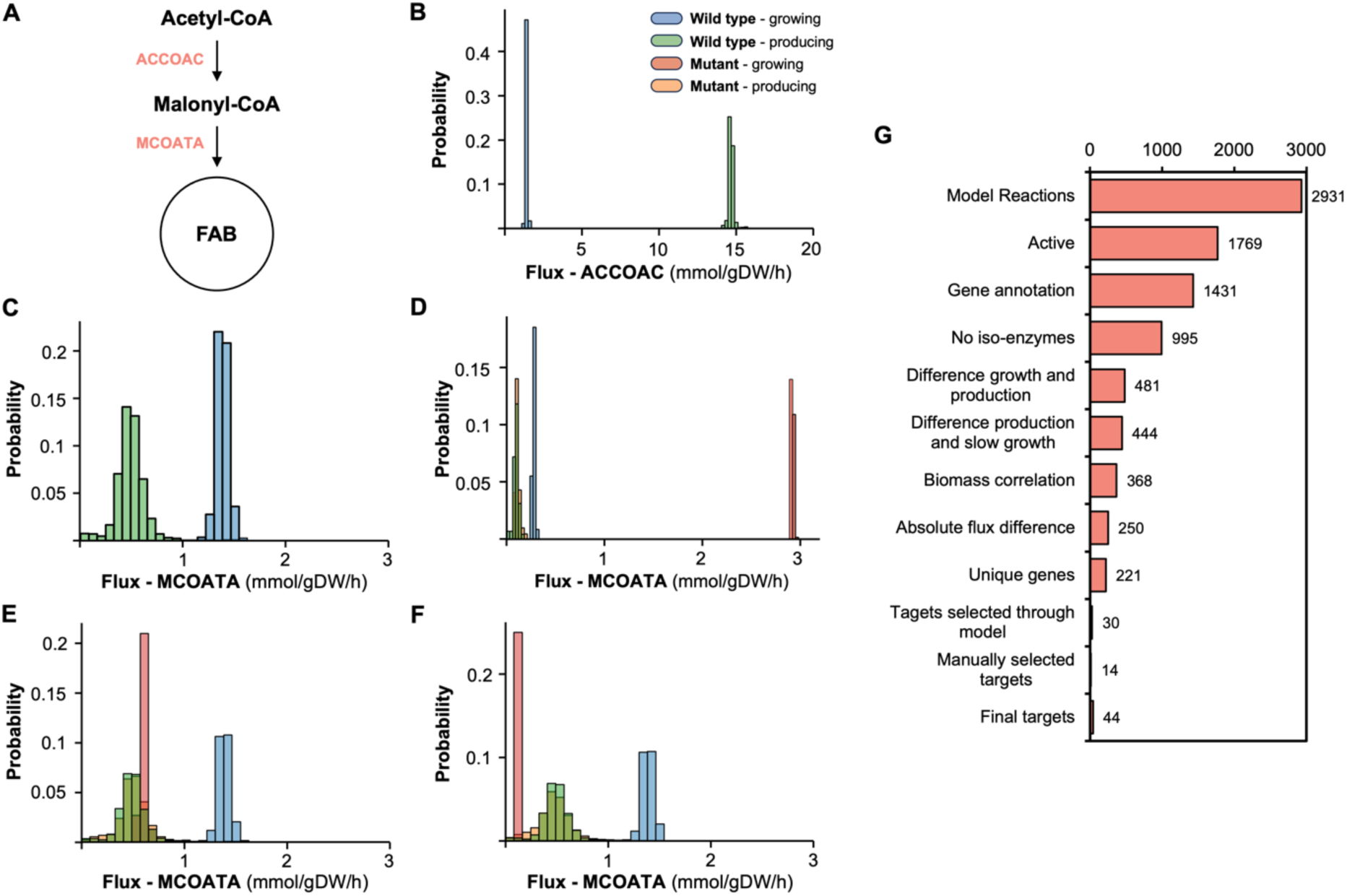
*In silico* prediction of gene targets through flux sampling. A) Schematic overview of two potential targets for intervention: acetyl-CoA carboxylase (ACCOAC) and malonyl-CoA-ACP transacylase (MCOATA). B) ACCOAC shows low flux during growth (blue) while flux during production is high (green) and is thus a suitable target for overexpression. C) MCOATA shows high flux during growth and decreased flux during production making it a suitable knockdown target. D) In simulations of the ACC overexpression strain, the MCOATA flux during growth is increased compared to the wild-type, while the flux during production is similar, indicating that the effect of the knockdown is potentially higher (synergy). In comparison, the *lysC* knockdown strain E) shows a smaller difference between growth and production flux, indicating that the MCOATA knockdown is potentially less effective in this strain (anti-synergy). In the *tyrB* knockdown strain F), the flux through MCOATA is now lower instead of higher in the growth condition, switching the MCOATA prediction from a potential target for knock-down to overexpression. G) Overview of the number of targets remaining after applying subsequent filtering criteria. Note that the final number of genes target is higher than the number of strains tested, as some of the genes are situated in operons and are targeted together.

#### 3.1.2 Screening of selected gene knockdowns

A CRISPRi library was constructed to target each of the 44 selected genes, enabling dCas9-based downregulation. To rapidly evaluate the knockdown targets, we utilized a recently developed colorimetric malonyl-CoA biosensor (Yang et al., 2018) based on the tetrahydroxynaphthalene synthase (RppA) of *Streptomyces coelicolor*. This enzyme catalyzes the conversion of five malonyl-CoA molecules into one molecule of 1,3,6,8-tetrahydroxynaphthalene (THN) which is then spontaneously converted into the red-colored flaviolin in the presence of oxygen. This system was adapted to fulfil the needs of the current workflow resulting in the biosensor plasmid pFLAV (see supplementary materials). The complete CRISPRi library was introduced into *P. putida* carrying pFLAV, and the knockdown efficiency was assessed based on the relative flaviolin signal (Fig. 3A). Among the 44 screened gene targets, 11 exhibited more than a 20% increase in the relative fla-violin signal compared to the control strain (Fig. 3A). As expected, but also as predicted, the top five knockdown targets were related to fatty acid biosynthesis (FAB) (Fig. 3 A, B). The highest flaviolin production was achieved by downregulating the *fabD* gene, encoding malonyl-CoA-ACP transacylase, which catalyzes the first reaction of the fatty acid biosynthesis (Fig. 3B). Relative production increased by 5.4-fold, highlighting the importance of inhibiting fatty acid biosynthesis in malonyl-CoA availability. It is worth noting that along with the downregulation of *fabD* there is a simultaneous downregulation of *fabG* (3-ketoacyl-ACP reductase) (Fig. 3B) since both genes are in the same operon with *fabD* being upstream to *fabG*. In addition, interfering with *fabV*, *fabA*, *fabF* and *fabB* (Fig. 3A) showed a relative increase of 3-, 2.6-, 2.5-and 2.5-fold, respectively. In-terestingly, targeting the TCA cycle related enzymes did not appear to increase malonyl-CoA availability as initially expected. The only exception was *sucAB* (succinyl-CoA synthase), the re-pression of which increased flaviolin production by 1.36-fold. The results above indicate that inhi-bition of the downstream malonyl-CoA metabolism significantly improves its availability, whereas that was not found to apply for the upstream metabolism.

**Figure 3.**
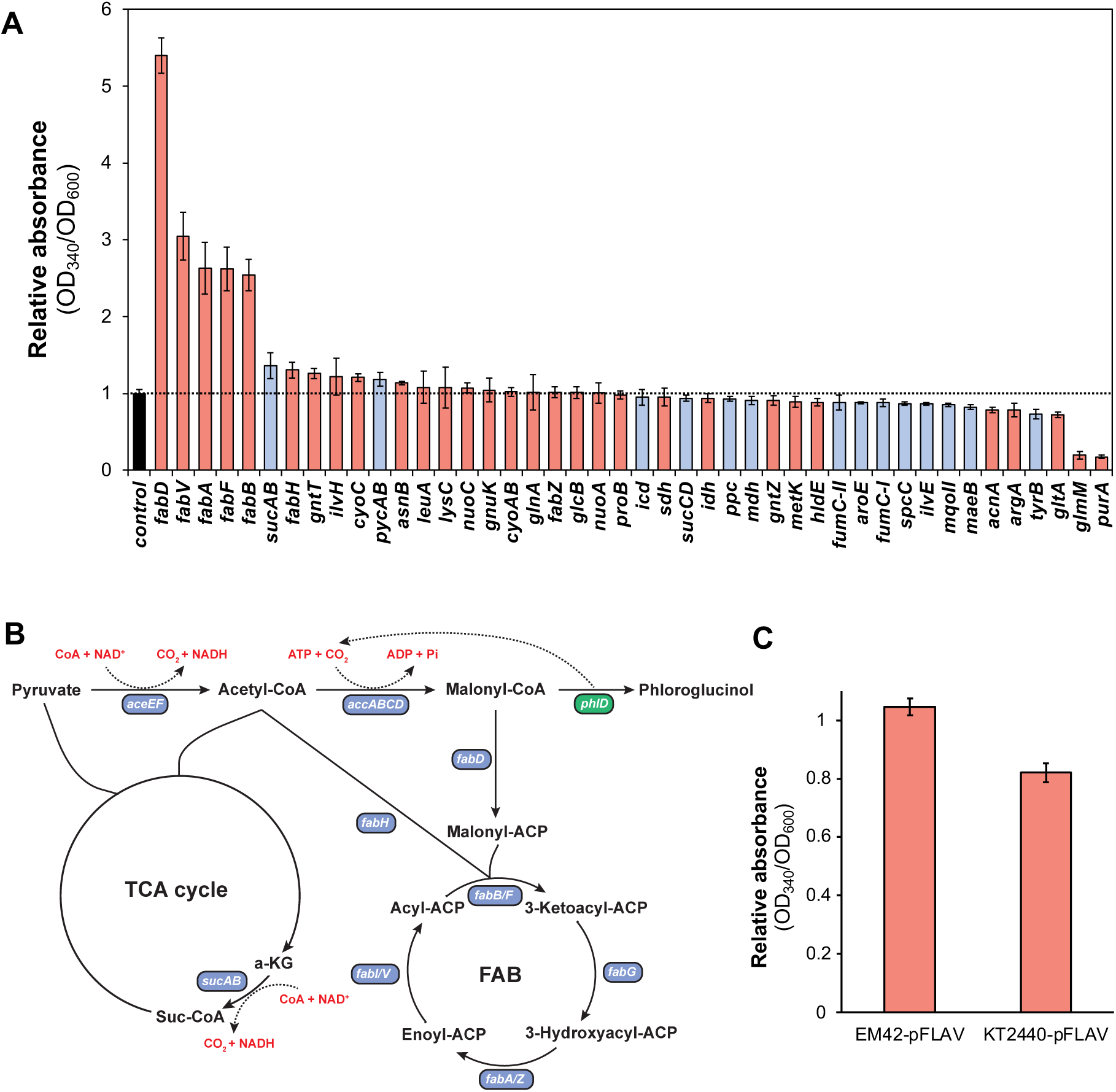
A) Biosensor mediated screening of gene knockdowns for increased malonyl-CoA availability. Relative flaviolin production from strains carrying pFLAV and each of the 44 CRISPRi target plasmids. Red bars; predicted gene targets, blue bars; rationally selected gene targets. B) Graphical representation of metabolic pathways associated with malonyl-CoA and phloroglucinol metabolism. Native genes are represented in blue boxes and heterologous in green. C) Relative flaviolin production by *P. putida* KT2440 and *P. putida* EM42 in MOPS minimal medium with 70 mM of glycose after 24h. Error bars indicate standard deviation of the data (n=3).

### 3.2 LEVEL 2: chassis selection

In bacteria, the conversion of acetyl-CoA to malonyl-CoA is catalyzed by the ATP-dependent ac-etyl-CoA carboxylase complex, AccABCD (ACC) (Fig. 3B). Both the overexpression of ACC and enhanced ATP supply have been shown to increase malonyl-CoA levels in *E. coli* (Yang et al., 2018; Tokuyama et al., 2019). To ensure that enough ATP would be available to facilitate efficient conversion of acetyl-CoA to malonyl-CoA, we decided to employ the streamlined *P. putida* EM42 strain. *P. putida* EM42 is a genome-reduced variant in which, among others, the flagellar machinery has been deleted, increasing ATP and NAD(P)H availability in the cell (Martínez-García et al., 2014; Lieder et al., 2015). As such, if ATP is a limiting factor, *P. putida* EM42 should exhibit more malonyl-CoA production than its parental strain. Both strains were transformed with pFLAV, grown in MOPS minimal medium and the relative flaviolin absorbance was measured after 24 h. The strain *P. putida* EM42 showed greater performance on flaviolin production, resulting in a 1.27-fold increase compared to *P. putida* KT2440 (Fig. 3C). Thus, we utilized *P. putida* EM42 as the base strain for further engineering.

### 3.3 LEVEL 3: RBS optimization of the ACC complex

To enhance the conversion rate of acetyl-CoA to malonyl-CoA, we focused on optimizing the expression of genes encoding the ACC subunits, *accABCD*, by RBS engineering. Using a deep learning model for RBS prediction (Höllerer et al., 2020) in combination with the RBS library opti-mization algorithm RedLibs^42^, we generated, *in silico*, six RBS variants per gene that uniformly cover the accessible range of predicted translation initiation rates resulting in a total of 1296 RBS combinations (Fig. 4A). We then constructed all combinations in a synthetic *accABCD* operon library, which was subsequently integrated directly into *P. putida*’s chromosome. To do this, we developed a genetic strategy tailored for Gram-negative bacteria that enables highly efficient chromosomal integration of large operons. The strategy relies on Cre-mediated recombination and follows the concept of the genome integration method CRAGE (Wang et al., 2019). At first, we genomically integrated into the chromosome of *P. putida* EM42 a *cre* transcriptional unit (TU) using the pGNW suicide vector (Wirth et al., 2019). The *cre* coding sequence along with its RBS was flanked by the *lox71* and *lox2/66* sites, whereas its expression was driven by the constitutive promoter BBa_J23100 and terminated by the T7 terminator (Amarelle et al., 2019) (Fig. 4B). In this setup, after recombination, *cre* is replaced by the insert, while the promoter and terminator remain in place to control the expression of the inserted operon of interest. To transfer the inte-gration cassette, we constructed a suicide vector called pLGR. pLGR consists of the promoter-less insert of interest and the antibiotic marker *acc* (resistance to gentamycin) flanked by *lox* sites (Fig. 4B). When the pLGR plasmid is transformed into the recipient strain, the insert and the antibiotic marker will replace the genomically integrated *cre* (Fig. 4B) allowing the selection of the success-fully engineered cells using the corresponding antibiotic. The high efficiency of the method was confirmed by integrating inserts of various sizes (see supplementary materials).

**Figure 4.**
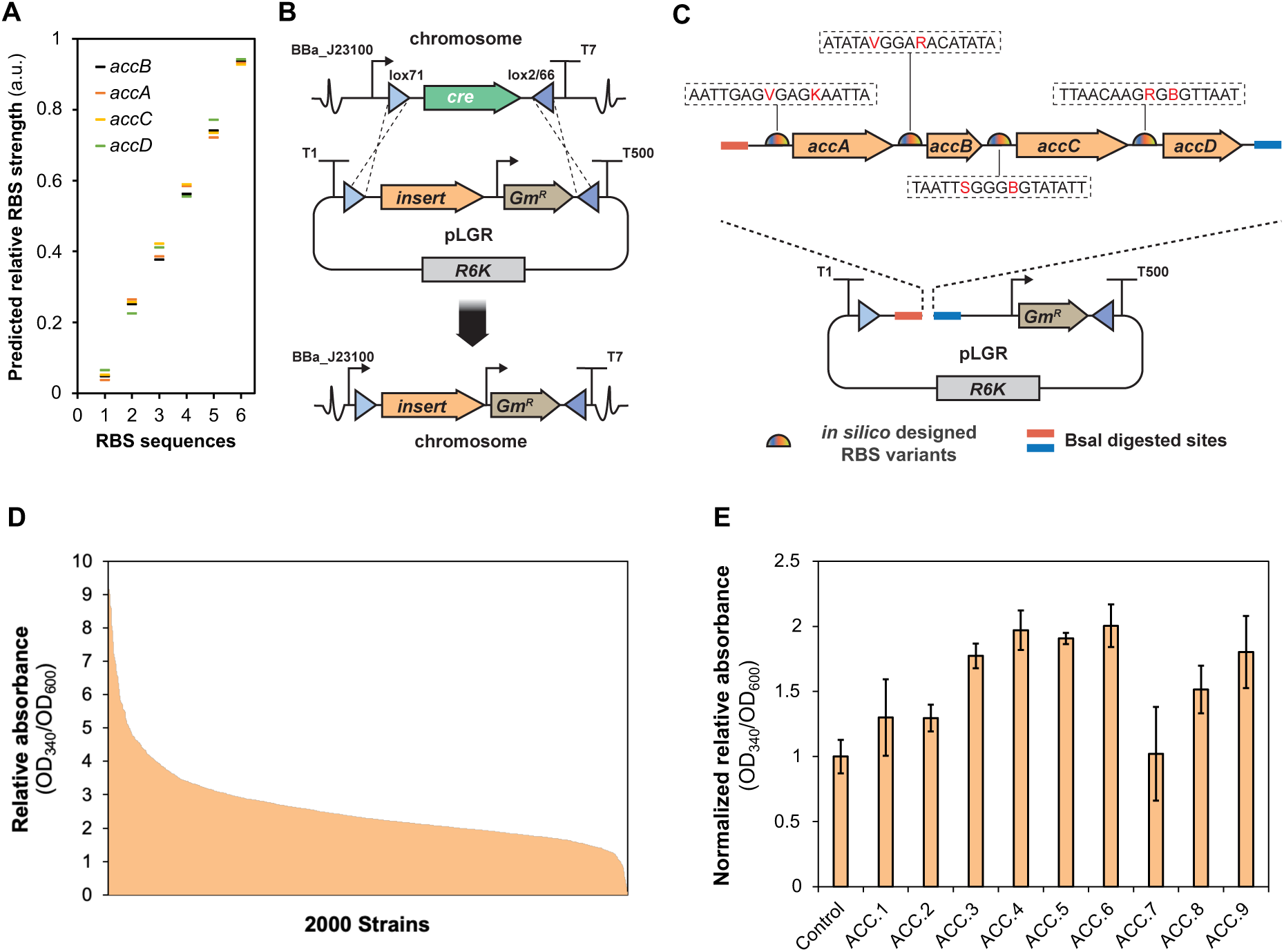
High-throughput genome integration of the combinatorial *accABCD* RBS library. A) Predicted relative RBS strength of each candidate RBS sequence for *accA*, *accB*, *accC* and *accD*. B) Graphical representation of the CRAGE-like recombination method that allows high-throughput genome inte-gration into *P. putida*’s genome. At first, a Cre-*lox* cassette is integrated into the chromosomal locus of interest using the suicide vector pGNW. The cassette comprised the Cre recombinase gene flanked by two *lox* sites, while the constitutive promoter BBa_J23100 was placed upstream the *lox71* and the transcriptional terminator T7 was placed downstream *lox2/66*. Later, a second suicide plasmid, pLGR, carrying the insert of interest and a gentamycin resistant gene TU (*acc*) flanked by two *lox* sites is electroporated into the recipient strain carrying the genome integrated Cre cassette. Integration is mediated through Cre-*lox* recombination. C) An *accABCD* operon library carrying all predicted RBS combinations was created using degenerate oligos and cloned into the vector pLGR through Golden Gate. D) Relative flaviolin production of 2000 *P. putida* EM42 mutants carrying the combinatorial *ac-cABCD* RBS library grown in LB medium. E) Relative flaviolin production from mutants integrated with the top 9 *accABCD* operons grown in MOPS medium with glucose. Error bars indicate standard devi-ation of the data (n=3).

The RBS library was constructed by PCR amplification of each *acc* gene using degenerate oligo-nucleotides coding for the six RBS options per gene, and the PCR products were subsequently cloned via Golden Gate into pLGR (Fig. 4C) (detailed cloning procedure is described in Materials and Methods). Following the transformation of *E. coli*, 40 colonies were PCR amplified and, after overnight growth, ten plasmids were extracted and sequenced. To our surprise, although 35 out of 40 colonies showed the correct PCR size, only four out of ten were shown to be correct after sequencing. Noteworthy, all sequenced colonies showed the expected PCR size before extraction indicating that plasmids were mutated during the overnight growth. All mutated plasmids lacked the complete sequence of *accABC* genes while a small fragment of *accD* remained in place. This observation strongly indicates the toxic effect caused by the *accABCD* operon even when promoter-less. Presumably, the constitutive promoters responsible for expressing the functional plasmid elements expressed the operon to a sufficient extent. To address this issue, we trans-formed the assembled library directly into *P. putida* EM42 carrying pFLAV and the genomically integrated Cre-*lox* cassette. The direct transformation was equally effective when mutants were plated on LB agar plates with gentamicin and kanamycin, resulting in a plurality of colonies. To test the integration efficiency, 40 colonies were PCR amplified, with 36 colonies displaying the correct insert size even after overnight growth. Having confirmed the high integration rate, a total of 2000 colonies were picked and inoculated into deep 96-well plates containing 200 μl of LB with the same antibiotics. Cells were grown overnight and flaviolin production was analyzed, as de-scribed above, to identify the RBS combinations that led to increased malonyl-CoA production (Fig. 4D). From these data, the top nine flaviolin producers were selected and cultivated in 14 ml falcon tubes with MOPS minimal medium supplemented with 70 mM glucose. Seven of them showed significantly increased signal compared to the control (Fig. 4E), with the greatest achiev-ing a 2-fold increase compared to the control strain (EM42 transformed with an empty pLGR vec-tor). Notably, the final cell density was not affected for any of the mutants analyzed. The best producing clone (ACC.6) was sequenced to identify the optimal RBS combination (see supple-mentary materials). Following, we reconstructed the complete transcriptional unit excluding the gentamicin resistant gene and integrated it into *P. putida* EM42 genome using the suicide vector pGNW. The new marker-less *P. putida* EM42^ACCopt^ was utilized for further experimentation.

### 3.4 Combinatorial engineering for enhancing phloroglucinol production

The primary purpose of increasing malonyl-CoA levels is to optimize other biosynthetic pathways of interest. To this end, we selected phloroglucinol, a polyketide with diverse applications in the pharmaceutical and chemical industries, as our target product. Phloroglucinol has antibacterial and antiviral properties and serves as a precursor for cosmetics, sedatives, and explosives (Yang et al., 2012). Using the *P. fluorescens* phloroglucinol synthase, encoded by the *phlD* gene, various organisms, including *E. coli* and *Arabidopsis*, have been engineered towards the sustainable production of phloroglucinol (Yang et al., 2012; Abdel-Ghany et al., 2016). To construct a *P. putida* strain capable of producing phloroglucinol, the codon-optimized *P. fluorescens* Pf-5 *phlD* was cloned into vector pSEVAb25 under the control of the RhaRS/*P_rhaBAD_* inducible expression system resulting in plasmid pPhlo. Shake flask phloroglucinol production experiments in MOPS minimal medium with glucose were performed for six *P. putida* strains with different genotypes selected based on the RppA-biosensor screening.

The EM42 strain carrying pPhlo and a non-targeting CRISPRi vector successfully produced 0.26 mM of phloroglucinol. ACC optimization (strain EM42^ACCopt^ carrying pPhlo and a non-targeting CRISPRi vector) increased production titer by 2.1-fold, reaching 0.56 mM after 24 hours (Fig. 5C). For the downregulation targets, we selected *sucAB* and *fabD*. *SucAB* demonstrated to be the most effective target among the genes related to upstream metabolism, and similarly *fabD* from the downstream malonyl-CoA metabolism. Both CRISPRi vectors were individually transformed in strain EM42^ACCopt^ carrying pPhlo, resulting in EM42^ACCopt^_*sucAB* and EM42^ACCopt^_*fabD*, respectively. The downregulation of *sucAB* and *fabD* resulted in significant production boosts, with phloroglucinol levels increasing by 2.92-fold and 4.60-fold, respectively (Fig. 5C). Given that *fabD* downregulation inhibits malonyl-CoA degradation and *sucAB* downregulation possibly increases the acetyl-CoA pool, we hypothesized a synergistic effect from their combined downregulation. Indeed, the double downregulation of *fabD* and *sucAB* led to a significant 5.8-fold increase in phloroglucinol titer achieving 1.53 mM after 24 h (Fig. 5C). Furthermore, the final cell densities were slightly decreased, despite an increase in glucose consumption (Fig. 5A, B). This indicates a CRISPRi-mediated flux redirection from growth to product synthesis. These results demonstrate that simultaneous overexpression of the ACC complex and inhibition of both the upstream and downstream metabolism of malonyl-CoA synergistically enhance its intracellular pool and consequently phloroglucinol production.

**Figure 5.**
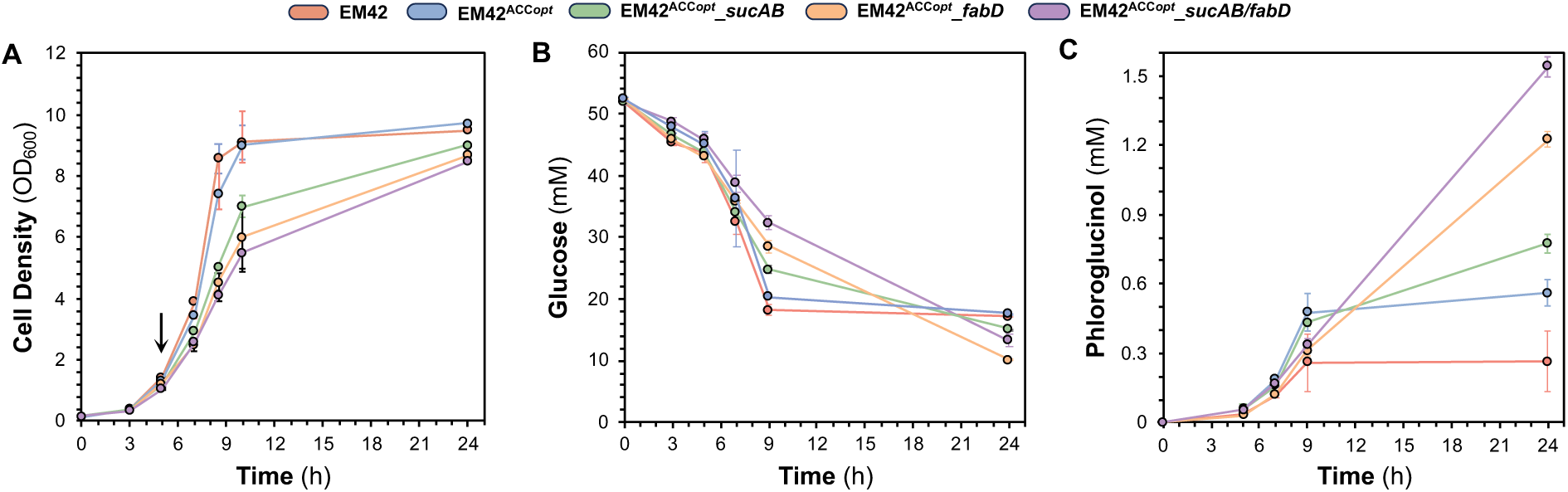
Phloroglucinol production cultivation data from *P. putida* mutants with various genotypes. A) Growth profile, B) glucose consumption, and C) phloroglucinol titer from selected mutants growing in minimal medium with glucose. The arrow in panel A indicates when the inducer L-Rhamnose was added. Error bars indicate standard deviation of the data (n=3).

## 4. DISCUSSION

In this study, by making use of modern metabolic engineering technologies, we employed a multi-level engineering workflow to enhance intracellular malonyl-CoA levels in *P. putida*, which has substantial potential as industrial chassis. This workflow integrates computational, genetic, and screening tools to systematically explore various metabolic targets, facilitating rapid achievement of our overproduction objectives.

In the first optimization level, we focused on the *in silico* identification of knockdown targets that show significant flux modulation when comparing growth versus malonyl-CoA producing phenotypes. Here flux sampling is used to explore the whole solution space of feasible fluxes compatible with maximum growth or production instead (Megchelenbrink et al., 2014). Arguably, this can be more representative, for example if the optimal flux profile is not unique, or significantly different from the average flux profile. In contrast to other strain design methods, which are often based on the design of a single strain with multiple knockouts to achieve growth-coupling (Burgard et al., 2003), our method has the advantage that reaction targets can be predicted independently, and do not require the implementation of several knockouts at once. The method here presented can be further extended with additional selection criteria (Zhou et al., 2020). These properties make it well suited for the high-throughput parallel screening enabled through a CRISPRi library and the malonyl-CoA responsive biosensor.

Experimental evaluation of a limited set of gene knockdowns, encompassing central metabolism, revealed a strong positive effect from inhibiting downstream malonyl-CoA metabolism. Specifi-cally, interfering with *fabD*, the first reaction of FAB, led to a 5.4-fold increase in the relative bio-sensor signal. This is the first study to demonstrate the inhibition of *fabD* in *P. putida*, guiding future research related to the malonyl-CoA node. Conversely, interfering with upstream metabo-lism had minimal impact on the malonyl-CoA pool, suggesting that acetyl-CoA is not a bottleneck.

In parallel with the identification of knockdown targets, we explored the potential of increasing the conversion rate of acetyl-CoA to malonyl-CoA. Previous studies in other organisms have shown that the overexpression of ACC significantly improved the intracellular malonyl-CoA pool resulting in high production rates of several malonyl-CoA-derived products (Davis et al., 2000; Zhou et al., 2020). However, overexpression of this enzymatic complex often imposes a metabolic burden on cells, which is commercially undesirable (Coussement et al., 2014). We therefore fo-cused on harmonizing the expression levels of all ACC subunits, instead of directly overexpressing the entire complex. In addition, since the production of malonyl-CoA is an ATP-dependent pro-cess, we employed the streamlined EM42 strain, which, compared to KT2440, showed enhanced ability to produce flaviolin, presumably due to its elevated ATP levels. To coordinate the abun-dance of the four enzymes of the ACC complex, we designed a library of RBS sequences for all *accABCD* genes. Given that the RppA-based colorimetric screening is slow, the number of the potential RBS combinations had to be limited to a manageable extent. To overcome this limita-tion, we utilized the RBS prediction and smart library design (RedLibs algorithm) to construct a library that meets the throughput of the RppA-biosensor and, at the same time, spans the entire translation initiation rate space. After constructing and screening thousands of variants of a syn-thetic *accABCD* operon, we identified RBS combinations that enabled increased malonyl-CoA production up to 2-fold without affecting cell growth.

An important aspect of our RBS optimization approach was the integration of the entire operon library directly into the genome of *P. putida* using a customized Cre-mediated integration system. In comparison, since the required mutants usually range from hundreds to thousands, these efforts often rely on plasmid-based expression systems (Rennig et al., 2019; Su et al., 2020). However, as the genetic background of any plasmid-based system differs significantly from that of a chromo-somal integration, the optimization effort would be wasted when the system has to eventually be integrated genomically to ensure genetic stability and reduce metabolic burden.

The identified most-promising genetic interventions were applied in combination to enhance the production of a representative proof-of-concept malonyl-CoA product, phloroglucinol, achieving a significant 5.8-fold increase on titer. Noteworthy, the maximum titer was a result of the *sucAB/fabD* double downregulation in the ACC optimized strain. Despite the beneficial effect of *sucAB* downregulation, it was predicted as a target for overexpression instead during the initial *in silico* analysis. Upon further investigation of this mismatch, we found that the modeled flux through the reactions catalyzed by succinyl-CoA synthetase was highly variable in the production case due to loops in the model, resulting in an unrealistically high mean flux. After correcting for this effect, by combining the fluxes involved in this loop, it still resulted in a minor increase in absolute flux in the production scenario compared to the growth scenario, although with a change in overall reaction directionality. If the reaction is not reversible, as assumed in the model, this would indeed match the observed experimental data. This highlights that the accuracy of the model is essential for reliable productions, as incorrectly labeled reaction reversibilities can thus lead to mispredictions.

Using *in silico* predictions, we identified several targets that increased the relative flaviolin signal, though not all were successful. False positives are expected as flux sampling does not establish causality between targets and phenotypes, nor account for enzyme expression, thermodynamic favorability, or precursor availability. Improvements could involve refining selection criteria or sim-ulating additional conditions, such as increased co-factor availability, to better differentiate pro-duction pathway limitations. Refining growth and production scenarios with intermediate steps could further distinguish unconstrained reactions from those correlated with increased growth or production.

While we successfully enhanced malonyl-CoA levels, further improvements to the workflow are possible. Running the full optimization LEVEL1 using the EM42 strain might yield different knock-down targets due to its metabolic differences from the parental strain. However, a calibrated model for this strain is currently unavailable, so we used the wild-type strain KT2440. Furthermore, new recombineering tools in *P. putida* offer opportunities to interfere genomically with identified targets, increasing stability and enhancing industrial applicability (Asin-Garcia et al., 2021; Asin-Garcia et al., 2023). For example, ReScribe could facilitate the high-throughput exchange of native regulatory elements with synthetic counterparts, leading to genomically stable phenotypes (Asin-Garcia et al., 2021).

## FUNDING

We acknowledge the financial support by the European Union Horizon2020 projects EmPower-Putida (grant number 635536), IBISBA (grant numbers 730976 and 871118), BIOS (grant number 101070281) and Bioindustry 4.0 (grant number 101094287).

## DECLARATION OF COMPETING INTEREST

The authors declare no conflict of interest.

## DATA AVAILABILITY

Data will be made available on request.

## Notes

### Competing Interest Statement

The authors have declared no competing interest.

## REFERENCES

Abdel-Ghany, S. E., Day, I., Heuberger, A. L., Broeckling, C. D., Reddy, A. S., 2016. Production of Phloroglucinol, a Platform Chemical, in Arabidopsis using a Bacterial Gene. Sci Rep. 6, 38483. 10.1038/srep38483.

Amarelle, V., Sanches-Medeiros, A., Silva-Rocha, R., Guazzaroni, M. E., 2019. Expanding the Toolbox of Broad Host-Range Transcriptional Terminators for Proteobacteria through Metagenomics. ACS Synth Biol. 8(4), 647–654. 10.1021/acssynbio.8b00507.

Asin-Garcia, E., Garcia-Morales, L., Bartholet, T., Liang, Z., Isaacs, F. J., Martins Dos Santos, V. A. P., 2023. Metagenomics harvested genus-specific single-stranded DNA-annealing proteins improve and expand recombineering in Pseudomonas species. Nucleic Acids Res. 51(22), 12522– 12536. 10.1093/nar/gkad102.

Asin-Garcia, E., Martin-Pascual, M., Garcia-Morales, L., van Kranenburg, R., Martins Dos Santos, V. A. P., 2021. ReScribe: An Unrestrained Tool Combining Multiplex Recombineering and Min-imal-PAM ScCas9 for Genome Recoding *Pseudomonas putida*. ACS Synth. Biol. 10(10), 2672– 2688. 10.1021/acssynbio.1c00297.

Bagdasarian, M., Lurz, R., Rückert, B., Franklin, F. C., Bagdasarian, M. M., Frey, J., Timmis, K. N., 1981. Specific-purpose plasmid cloning vectors. II. Broad host range, high copy number, RSF1010-derived vectors, and a host-vector system for gene cloning in Pseudomonas. Gene, 16(1-3), 237–247. 10.1016/0378-1119(81)90080-9.

Batianis, C., Kozaeva, E., Damalas, S. G., Martín-Pascual, M., Volke, D. C., Nikel, P. I., Martins Dos Santos, V. A. P., 2020. An expanded CRISPRi toolbox for tunable control of gene expression in *Pseudomonas putida*. Microb. Biotechn. 13(2), 368–385. 10.1111/1751-7915.13533.

Bordbar, A., Monk, J. M., King, Z. A., Palsson, B. O., 2014. Constraint-based models predict metabolic and associated cellular functions. Nat. Rev. Genet. 15(2), 107–120. 10.1038/nrg3643.

Burgard, A. P., Pharkya, P., Maranas, C. D., 2003. Optknock: a bilevel programming framework for identifying gene knockout strategies for microbial strain optimization. Biotechnol. Bioeng. 84(6), 647–657. 10.1002/bit.10803.

Choi, K. H., Kumar, A., Schweizer, H. P., 2006. A 10-min method for preparation of highly electrocompetent *Pseudomonas aeruginosa* cells: application for DNA fragment transfer between chromosomes and plasmid transformation. J. Microbiol. Methods. 64(3), 391–397. 10.1016/j.mimet.2005.06.001.

Coussement, P., Maertens, J., Beauprez, J., Van Bellegem, W., De Mey, M., 2014. One step DNA assembly for combinatorial metabolic engineering. Metab. Eng. 23, 70–77. 10.1016/j.ymben.2014.02.012.

Dahiya, S., Katakojwala, R., Ramakrishna, S., Mohan, S. V., 2020. Biobased products and life cycle assessment in the context of circular economy and sustainability. Mater Circ Econ 2, 7. 10.1007/s42824-020-00007-x.

Damalas, S. G., Batianis, C., Martin-Pascual, M., de Lorenzo, V., Martins Dos Santos, V. A. P., 2020. SEVA 3.1: enabling interoperability of DNA assembly among the SEVA, BioBricks and Type IIS restriction enzyme standards. Microb. Biotechn. 13(6), 1793–1806. 10.1111/1751-7915.13609.

Davis, M. S., Solbiati, J., Cronan, J. E., Jr., 2000. Overproduction of acetyl-CoA carboxylase activity increases the rate of fatty acid biosynthesis in *Escherichia coli*. J. Biol. Chem. 275(37), 28593–28598. 10.1074/jbc.M004756200.

Dinh C.V., Prather K.L.J., 2019. Development of an autonomous and bifunctional quorum-sensing circuit for metabolic flux control in engineered *Escherichia coli*. Proc. Natl. Acad. Sci. U S A. 116(51), 25562–25568. doi:10.1073/pnas.1911144116.

Dos Santos, V. A., Heim, S., Moore, E. R., Strätz, M., Timmis, K. N., 2004. Insights into the genomic basis of niche specificity of *Pseudomonas putida* KT2440. Environ. Microbiol. 6(12), 1264– 1286. 10.1111/j.1462-2920.2004.00734.x.

Ebrahim, A., Lerman, J. A., Palsson, B. O., Hyduke, D. R., 2013. COBRApy: COnstraints-Based Reconstruction and Analysis for Python. BMC Syst. Biol. 7, 74. 10.1186/1752-0509-7-74.

Ferreira, R., Skrekas, C., Hedin, A., Sánchez, B. J., Siewers, V., Nielsen, J., David, F., 2019. Model-Assisted Fine-Tuning of Central Carbon Metabolism in Yeast through dCas9-Based Regulation. ACS Synth. Biol. 8(11), 2457–2463. 10.1021/acssynbio.9b00258

Geweke, J. F., 1991. Evaluating the accuracy of sampling-based approaches to the calculation of posterior moments. Staff Report 148.

Gläser, L., Kuhl, M., Jovanovic, S., Fritz, M., Vögeli, B., Erb, T. J., Becker, J., Wittmann, C., 2020. A common approach for absolute quantification of short chain CoA thioesters in prokaryotic and eukaryotic microbes. Microb. Cell Fact. 19(1), 160. 10.1186/s12934-020-01413-1.

Green, R., Rogers, E. J., 2013. Transformation of chemically competent *E. coli*. Methods Enzymol. 529, 329–336. 10.1016/B978-0-12-418687-3.00028-8.

Hanahan D., 1983. Studies on transformation of *Escherichia coli* with plasmids. J. Mol. Biol. 166(4), 557–580. 10.1016/s0022-2836(83)80284-8.

Höllerer, S., Papaxanthos, L., Gumpinger, A. C., Fischer, K., Beisel, C., Borgwardt, K., Benenson, Y., Jeschek, M., 2020. Large-scale DNA-based phenotypic recording and deep learning enable highly accurate sequence-function mapping. Nat. Commun. 11(1), 3551. 10.1038/s41467-020-17222-4.

Jeschek, M., Gerngross, D., Panke, S., 2016. Rationally reduced libraries for combinatorial path-way optimization minimizing experimental effort. Nat. Commun. 7, 11163. 10.1038/ncomms11163.

Johnson, A. O., Gonzalez-Villanueva, M., Wong, L., Steinbüchel, A., Tee, K. L., Xu, P., Wong, T. S., 2017 Design and application of genetically-encoded malonyl-CoA biosensors for metabolic engineering of microbial cell factories. Metab. Eng. 44, 253–264. doi:10.1016/j.ymben.2017.10.01.

Kampers, L. F. C., Volkers, R. J. M., Martins Dos Santos, V. A. P., 2019. *Pseudomonas putida* KT2440 is HV1 certified, not GRAS. Microb. Biotechnol. 12(5), 845–848. 10.1111/1751-7915.13443.

Ko, Y. S., Kim, J. W., Lee, J. A., Han, T., Kim, G. B., Park, J. E., Lee, S. Y., 2020. Tools and strategies of systems metabolic engineering for the development of microbial cell factories for chemical production. Chem Soc Rev. 49(14), 4615–4636. doi:10.1039/d0cs00155d.

LaBauve, A. E., & Wargo, M. J., 2012. Growth and laboratory maintenance of *Pseudomonas ae-ruginosa*. Curr. Protoc. Microbiol. Chapter 6, Unit–6E.1. 10.1002/9780471729259.mc06e01s25.

Lewis, N. E., Hixson, K. K., Conrad, T. M., Lerman, J. A., Charusanti, P., Polpitiya, A. D., Adkins, J. N., Schramm, G., Purvine, S. O., Lopez-Ferrer, D., Weitz, K. K., Eils, R., König, R., Smith, R. D., Palsson, B. Ø., 2010. Omic data from evolved *E. coli* are consistent with computed optimal growth from genome-scale models. Mol. Syst. Biol. 6, 390. 10.1038/msb.2010.47.

Li, S., Li, Z., Pang, S., Xiang, W., Wang, W., 2021. Coordinating precursor supply for pharmaceutical polyketide production in Streptomyces. Curr. Opin. Biotechnol. 69, 26–34. 10.1016/j.copbio.2020.11.006

Lieder, S., Nikel, P. I., de Lorenzo, V., Takors, R., 2015. Genome reduction boosts heterologous gene expression in *Pseudomonas putida*. Microb. Cell Fact. 14, 23. 10.1186/s12934-015-0207-7.

Liu, R., Liang, L., Choudhury, A., Bassalo, M. C., Garst, A. D., Tarasava, K., Gill, R. T., 2018. Iterative genome editing of *Escherichia coli* for 3-hydroxypropionic acid production. Metab. Eng. 47, 303–313. 10.1016/j.ymben.2018.04.00711.

Liu, H., Jin, Y., Zhang, R., Ning, Y., Yu, Y., Xu, P., Deng, L., Wang, F., 2023. Recent advances and perspectives on production of value-added organic acids through metabolic engineering. Biotechnol. Adv. 62, 108076. 10.1016/j.biotechadv.2022.108076

Loeschcke, A., Thies, S., 2015. *Pseudomonas putida*-a versatile host for the production of natural products. Appl. Microbiol. Biotechnol. 99(15), 6197–6214. 10.1007/s00253-015-6745-4.

Lu, C., Batianis, C., Akwafo, E. O., Wijffels, R. H., Martins Dos Santos, V. A. P., Weusthuis, R. A., 2021. When metabolic prowess is too much of a good thing: how carbon catabolite repression and metabolic versatility impede production of esterified α,ω-diols in *Pseudomonas putida* KT2440. Biotechnol. Biofuels. 14(1), 218. 10.1186/s13068-021-02066-x.

Ma, J., Gu, Y., Marsafari, M., Xu, P., 2020. Synthetic biology, systems biology, and metabolic engineering of *Yarrowia lipolytica* toward a sustainable biorefinery platform. J. Ind. Microbiol. Biotechnol. 47(9-10), 845–862. 10.1007/s10295-020-02290-8.

Machado, D., Herrgård, M. J., 2015. Co-evolution of strain design methods based on flux balance and elementary mode analysis. Metab. Eng. Commun. 2, 85–92. 10.1016/j.meteno.2015.04.001.

Martin-Pascual, M., Batianis, C., Bruinsma, L., Asin-Garcia, E., Garcia-Morales, L., Weusthuis, R. A., van Kranenburg, R., Martins Dos Santos, V. A. P., 2021. A navigation guide of synthetic biology tools for *Pseudomonas putida*. Biotechnol. Adv. 49, 107732. 10.1016/j.biotechadv.2021.107732.

Martínez-García, E., Nikel, P. I., Aparicio, T., de Lorenzo, V., 2014. Pseudomonas 2.0: genetic upgrading of *P. putida* KT2440 as an enhanced host for heterologous gene expression. Microb. Cell Fact. 13, 159. 10.1186/s12934-014-0159-3.

Megchelenbrink, W., Huynen, M., Marchiori, E., 2014. optGpSampler: an improved tool for uniformly sampling the solution-space of genome-scale metabolic networks. PloS one. 9(2), e86587. 10.1371/journal.pone.0086587.

Mezzina, M. P., Manoli, M. T., Prieto, M. A., Nikel, P. I., 2021. Engineering Native and Synthetic Pathways in *Pseudomonas putida* for the Production of Tailored Polyhydroxyalkanoates. Biotechnol. J. 16(3), e2000165. 10.1002/biot.202000165.

Milke, L., Kallscheuer, N., Kappelmann, J., Marienhagen, J., 2019. Tailoring *Corynebacterium glutamicum* towards increased malonyl-CoA availability for efficient synthesis of the plant pentaketide noreugenin. Microb. Cell. Fact. 18(1), 71. 10.1186/s12934-019-1117-x.

Milke, L., Marienhagen, J., 2020. Engineering intracellular malonyl-CoA availability in microbial hosts and its impact on polyketide and fatty acid synthesis. Appl. Microbiol. Biotechnol. 104(14), 6057–6065. 10.1007/s00253-020-10643-7.

Moreno-Paz, S., van der Hoek, R., Eliana, E., Zwartjens, P., Gosiewska, S., Martins Dos Santos, V. A. P., Schmitz, J., Suarez-Diez, M., 2024. Machine Learning-Guided Optimization of p-Coumaric Acid Production in Yeast. ACS Synth. Biol. 13(4), 1312–1322. 10.1021/acssynbio.4c00035.

Nogales, J., Mueller, J., Gudmundsson, S., Canalejo, F. J., Duque, E., Monk, J., Feist, A. M., Ramos, J. L., Niu, W., Palsson, B. O., 2020. High-quality genome-scale metabolic modelling of *Pseudomonas putida* highlights its broad metabolic capabilities. Environ. Microbiol. 22(1), 255–269. 10.1111/1462-2920.14843.

Poblete-Castro, I., Becker, J., Dohnt, K., dos Santos, V. M., Wittmann, C., 2012. Industrial biotechnology of *Pseudomonas putida* and related species. Appl. Microbiol. Biotechnol. 93(6), 2279–2290. 10.1007/s00253-012-3928-0.

Raman, S., Rogers, J. K., Taylor, N. D., Church, G. M., 2014. Evolution-guided optimization of biosynthetic pathways. Proc. Natl. Acad. Sci. U S A. 111(50), 17803–17808. 10.1073/pnas.1409523111.

Rennig, M., Mundhada, H., Wordofa, G. G., Gerngross, D., Wulff, T., Worberg, A., Nielsen, A. T., Nørholm, M. H. H., 2019. Industrializing a Bacterial Strain for l-Serine Production through Translation Initiation Optimization. ACS Synth. Biol. 8(10), 2347–2358. 10.1021/acssynbio.9b00169.

Solovyev, V., Salamov, A., 2011. Automatic annotation of microbial genomes and metagenomic sequences. Metagenomics and its applications in agriculture. Biomedicine and Environmental studies 61–78.

Su, H. H., Peng, F., Ou, X. Y., Zeng, Y. J., Zong, M. H., Lou, W. Y., 2020. Combinatorial synthetic pathway fine-tuning and cofactor regeneration for metabolic engineering of *Escherichia coli* significantly improve production of D-glucaric acid. N. Biotechnol. 59, 51–58. 10.1016/j.nbt.2020.03.004.

Tan, Z., Yoon, J. M., Chowdhury, A., Burdick, K., Jarboe, L. R., Maranas, C. D., Shanks, J. V., 2018. Engineering of *E. coli* inherent fatty acid biosynthesis capacity to increase octanoic acid production. Biotechnol. Biofuels. 11, 87. 10.1186/s13068-018-1078-z.

Tarasava, K., Liu, R., Garst, A., Gill, R. T., 2018. Combinatorial pathway engineering using type I-E CRISPR interference. Biotechnol. Bioeng., 115(7), 1878–1883. 10.1002/bit.26589.

Tokuyama, K., Toya, Y., Matsuda, F., Cress, B. F., Koffas, M. A. G., Shimizu, H., 2019. Magnesium starvation improves production of malonyl-CoA-derived metabolites in *Escherichia coli*. Metab. Eng. 52, 215–223. 10.1016/j.ymben.2018.12.002

van Rosmalen, R. P., Moreno-Paz, S., Duman-Özdamar, Z. E., Suarez-Diez, M., 2024. CFSA: Comparative flux sampling analysis as a guide for strain design. Metab Eng Commun. 19, e00244. 10.1016/j.mec.2024.e00244.

Voigt C. A., 2020. Synthetic biology 2020-2030: six commercially-available products that are changing our world. Nat. Commun. 11(1), 6379. 10.1038/s41467-020-20122-2

Volke, D. C., Friis, L., Wirth, N. T., Turlin, J., Nikel, P. I., 2020. Synthetic control of plasmid replication enables target-and self-curing of vectors and expedites genome engineering of *Pseudomonas putida*. Metab. Eng. Commun. 10, e00126. 10.1016/j.mec.2020.e00126.

Wang, G., Zhao, Z., Ke, J., Engel, Y., Shi, Y.M., Robinson, D., Bingol, K., Zhang, Z., Bowen, B., Louie, K., Wang, B., Evans, R., Miyamoto, Y., Cheng, K., Kosina, S., De Raad, M., Silva, L., Luhrs, A., Lubbe, A., Hoyt, D.W., Francavilla, C., Otani, H., Deutsch, S., Washton, N.M., Rubin, E.M., Mouncey, N.J., Visel, A., Northen, T., Cheng, J.F., Bode, H.B., Yoshikuni, Y., 2019. CRAGE enables rapid activation of biosynthetic gene clusters in undomesticated bacteria. Nat. Microbiol. 4, 2498–2510. 10.1038/s41564-019-0573-8.

Wirth, N. T., Kozaeva, E., Nikel, P. I., 2020. Accelerated genome engineering of *Pseudomonas putida* by I-SceI-mediated recombination and CRISPR-Cas9 counterselection. Microb. Bio-techn. 13(1), 233–249. 10.1111/1751-7915.13396.

Xu, P., Ranganathan, S., Fowler, Z. L., Maranas, C. D., Koffas, M. A., 2011. Genome-scale meta-bolic network modeling results in minimal interventions that cooperatively force carbon flux towards malonyl-CoA. Metab. Eng. 13(5), 578–587. 10.1016/j.ymben.2011.06.008.

Yang, D., Kim, W. J., Yoo, S. M., Choi, J. H., Ha, S. H., Lee, M. H., Lee, S. Y., 2018. Repurposing type III polyketide synthase as a malonyl-CoA biosensor for metabolic engineering in bacteria. Proc. Natl. Acad. Sci. U S A. 115(40), 9835–9844. 10.1073/pnas.1808567115

Yang, F., Cao, Y., 2012. Biosynthesis of phloroglucinol compounds in microorganisms--review. Appl. Microbiol. Biotechnol. 93(2), 487–495. 10.1007/s00253-011-3712-6.

Zhou, S., Lama, S., Jiang, J., Sankaranarayanan, M., Park, S., 2020. Use of acetate for the produc-tion of 3-hydroxypropionic acid by metabolically-engineered *Pseudomonas denitrificans*. Bio-resour. Technol. 307, 123194. 10.1016/j.biortech.2020.123194.

Zhou, S., Yuan, S. F., Nair, P. H., Alper, H. S., Deng, Y., Zhou, J., 2021. Development of a growth coupled and multi-layered dynamic regulation network balancing malonyl-CoA node to en-hance (2S)-naringenin biosynthesis in *Escherichia coli*. Metab. Eng. 67, 41–52. 10.1016/j.ymben.2021.05.

