## Supplementary Materials for "Computer assisted multi-level optimization of malonyl-CoA availability in *Pseudomonas putida*"

**Table S1.** Primers used in this study

| **Primer name** | **Sequence** | **Purpose** |
| --- | --- | --- |
| 86 | CTGGATTCTCACCAATAAAAAACG | Plasmid sequencing |
| 87 | TCTAGGGCGGCGGAT | Plasmid sequencing |
| C16 | TTTAATTAATAGCTTTCGCTAAGGATG | Confirmation of gRNA |
| Rv Cas9.Ind | GGTCTCATTCCGTTTATTTTGTTATCATTCTATAGTATTAAGTATTG | Confirmation of gRNA |
| Ep | AGCTTTCGCTAAGGATGATTTCTGGTCTCGGAATTCGCG | SevaBrick amplification |
| Pp | ACCTTGCCCTTTTTTGCCGGTCTCGACTGCAGCG | SevaBrick amplification |
| Ev | CCGGGTCTCAATTCCAGAAATCATCCTTAGCGAAAGC | pSEVAb linearization |
| Pv | AGGTCTCACAGTCCGGCAAAAAAGGG | pSEVAb linearization |
| Sp | CCCTTTTTTGCCGGACTGCAGCGGCCGCTAGGTCTCCTAGTA | SevaBrick amplification |
| Xp | GGATGATTTCTGGAATTCGCGGCCGGTCTCTACTAGA | SevaBrick amplification |
| OP1-F_BBa_B0034_ | CGCTAAGGATGATTTCTGGAATTCGCAGGCCGCTTGGTCTCAACTAGAGAAAGAGGAGAAATACTAGATG | RBS-CDS_1_ amplification |
| EvBBa_J23100 | AGGTCTCCTAGTAGCTAGCACTGTACCTAGGACTGAGCTAGCCGTCAACTCTAGAAGCGGC | Promoter insertion |
| accA-F | ATGAACCCGAATTTCCTCG | Construction of *accABCD* library |
| accA-R | AGGTCTCATTAGAGGCCGTAGCTCATC |  |
| accA.RBS-F | AGGTCTCAACTAAATTGAGVGAGKAATTAATGAACCCGAATTTCCTCG |  |
| accB-F | ATGGATATCCGTAAAGTCAAG |  |
| accB-R1 | TCAAACGATGGTGAACAGC |  |
| accB-R2 | AGGTCTCATCAAACGATGGTGAACAGC |  |
| accB.RBS-F | AGGTCTCACTAAATATAVGGARACATATAATGGATATCCGTAAAGTCAAG |  |
| accC-F | ATGTCTGGGAAGCTCGAAAAAG |  |
| accC-R | AGGTCTCATCACTCCTGGTTGGCC |  |
| accC.RBS-F | AGGTCTCATTGATAATTSGGGBGTATATTATGTCTGGGAAGCTCGAAAAAG |  |
| accD-R1 | TCACGCGACGGCAGC |  |
| accD-R2 | AGGTCTCAACTGTCACGCGACGGCAGC |  |
| cusF.TS1-F | AGGTCTCTCCCGGACCCAAGGGTCAGTAG | Construction of *lox-cre-lox* pGNW |
| cusF.TS1-R | AGGTCTCTAGCTAGCACTGTACCTAGGACTGAGCTAGCCGTCAACAGGGCGATTCAGATC |  |
| lox_RBS_cre-F1 | TACCGTTCGTATAGCATACATTATACGAAGTTATAAAGAGGAGAAATACTAGATGTCC |  |
| lox_RBS_cre-F2 | AGGTCTCAAGCTTACCGTTCGTATAGCATAC |  |
| lox_RBS_cre-R1 | TACCGTTCGTATAAGAAACCATATACGAAGTTATTTAATCGCCATCTTCCAG |  |
| lox_RBS_cre-R2 | AGGTCTCAGAAGAGGAAGGTCGCGGCGAGCACGAGCGTACCGTTCGTATAAGAAACC |  |
| cusF.TS2-F | AGGTCTCACTTCATGCACCTCTTCGGCATCGGGCTGCACAAACTTCCTAGATGCTTCC |  |
| cusF.TS2-R | AGGTCTCATCGACAGCGTATTGGCTGG |  |
| RBS cre | AAAGAGGAGAAATACTAGATGTC |  |
| RBS cre-R | TTAATCGCCATCTTCCAG |  |
| R6K-F | AGGTCTCATTATTTCAGCTGCTGCCTGAG | Construction of pLGR |
| R6K-R | AGGTCTCATTATTTTCTCTTTGCGCTTGC |  |
| Gm+lox-F | AGGTCTCTATAACTTCGTATAGCATACATTATACGAACGGTAGGGGTCCCCAATAATTAC |  |
| Gm+lox-R | AGGTCTCTATAACTTCGTATAAGAAACCATATACGAACGGTATTAGGTGGCGGTACTTG |  |

**Table S2.** List of gRNA spacers used in this study

| **Spacer** | **Sequence (5’ - 3’)** |
| --- | --- |
| Non target | TGAGACCAGTCTCGGAAGCTCAAAGGTCTC |
| *fabA* | ACGGCAGCCTACAGGTAGACTATTGCGTTG |
| *fabB* | GATACGATGCCCAGACCAGTGATAACGACG |
| *fabD* | CTGAAAACTTGGATGGAAACACAGACCAAG |
| *fabF* | GCTCGCAAAGAAAAAACCGCACGCCAGCGA |
| *fabH* | AGGGGTTGTAGCCAGGGCTCGCCTGCGCAC |
| *fabV* | TTATCCGACACATATTACGGGGTGATCAGA |
| *fabZ* | GAGCCAGCTAGCGACTGTTCACTCTTGCTC |
| *gntT* | AAAACAACGATAAGACCGAGGCCTTCAAAC |
| *asnB* | GGCATCACCTCCCCAACGGAAAAACCGGTA |
| *cyoC* | AGTCAAGTAATGCACGGCGCTGCTCATGGTC |
| *leuA* | CATATCGTTATCGCTCGGCAGGCCCGATCT |
| *lysC* | ACGCCATTTCAATGGTGCCTCAGCCCATAC |
| *nuoC* | TTAAGAATGAAGCCAGGCCGCCGGCTCCTT |
| *gnuK* | GTTTGAAGGCCTCGGTCTTATCGTTGTTTT |
| *cyoAB* | TTGGTAGCCAACACTTCGTCCTGCCAAGTG |
| *glnA* | GGATAGTCGGCCAAGCTTACCTGCCTCACA |
| *glcB* | TCCGCGCCACGGGATACTACATGAAGCAGT |
| *nuoA* | AGTCGCCGCTCTACACTTATTTCAGACTGC |
| *proB* | CCGTGGATACAAAAACGCCGCTCCAGAGAG |
| *sdh* | TAGACAGTTAGGCTGCTTATGACAACGTGA |
| *idh* | GTATAGATGATCTTGGAACGGGTGGGCATG |
| *gntZ* | GCTTTTCAGATTAGTCCAGCATGCCGCCCG |
| *metK* | AGGTGAAAAGGGAGTATTCGCTCATCTCGA |
| *hldE* | AGGGGCAGGATATTAGCACAGGGTAGTCAA |
| *acnA* | TAGACCGGTTTACGCTACCTGACCTTTTTC |
| *argA* | GCAGGGGGCGTTCAGGATCATCGTCGGAAC |
| *glta* | AAATTGACATCTGAATTTATCCCTCTATAG |
| *glmM* | CCGTCGGTACCAAAGTATTTTCTGCTCATA |
| *purA* | ATTGGTCCTCATTCACGCAAACTTGGTTGC |
| *sucAB* | TTTCTTGCATGCTTGGTCACCCTCGATTAG |
| *icd* | CTGAGCCTAAACCACATCTGCCGACCTGCA |
| *sucCD* | CACGCAAGACTCACGACGGGCAGCCCGCCG |
| *mdh* | GTCGCATCGTCTTGGGTGGCCATTTAGCCC |
| *fumC-II* | TCACTCCTGTGTAGTTCGAAATCGCAGTTT |
| *fumC-I* | GATGAACTGATGCAAGGATTTGATGATGAG |
| *scpC* | GAGATTTCGCCCATTGCCGTCCCACGACCA |
| *ilvE* | TATGCGCACCATGCCACCTTAATACATCCT |
| *maeB* | GATACGCAAAGAGCCGGGGAGCCACAAGAC |
| *mqoII* | TGACGCTTGCAGCGACCGCCAGACCCAGCA |
| *tyrB* | CAAACGGGGAGTCAGTATAGTGATACCGAC |
| *pycAB* | TAGCCAGTCGCGTATTCACCCTAGCGCTGT |
| *ppc* | CCCCCAAGAGTGCCCACAGCCGAGCGACGA |
| *aroE* | GGTTACCAAAAACGACGTACTGGTCCATGA |

**Table S3.** Selection criteria for gene targets

| **Target** | **Selection criteria** |
| --- | --- |
| *fabA* | Predicted |
| *fabB* | Predicted |
| *fabD* | Predicted |
| *fabF* | Predicted |
| *fabH* | Predicted |
| *fabV* | Predicted |
| *fabZ* | Predicted |
| *gntT* | Predicted |
| *asnB* | Predicted |
| *cyoC* | Predicted |
| *leuA* | Predicted |
| *lysC* | Predicted |
| *nuoC* | Predicted |
| *gnuK* | Predicted |
| *cyoAB* | Predicted |
| *glnA* | Predicted |
| *glcB* | Predicted |
| *nuoA* | Predicted |
| *proB* | Predicted |
| *sdh* | Predicted |
| *idh* | Predicted |
| *gntZ* | Predicted |
| *metK* | Predicted |
| *hldE* | Predicted |
| *acnA* | Predicted |
| *argA* | Predicted |
| *glta* | Predicted |
| *glmM* | Predicted |
| *purA* | Predicted |
| *sucAB* | Rational selection; involved in TCA cycle, increased acetyl-CoA pool |
| *icd* |  |
| *sucCD* |  |
| *mdh* |  |
| *fumC-II* |  |
| *fumC-I* |  |
| *scpC* |  |
| *maeB* |  |
| *mqoII* |  |
| *ilvE* | Rational selection; save ATP |
| *tyrB* | Rational selection; save ATP |
| *pycAB* | Rational selection; save ATP |
| *ppc* | Rational selection; save ATP |
| *aroE* | Rational selection; save ATP |

**Table S4.** Best identified ACC RBS combination

| *accA* | AATTGAGGGAGGAATTA |
| --- | --- |
| *accB* | ATATAGGGAGACATATA |
| *accC* | TAATTGGGGTGTATATT |
| *accD* | TTAACAAGAGGGTTAAT |

Establishment of the RppA-biosensor in *P. putida*

The RppA-biosensor consists of a type III polyketide synthase, 1,3,6,8-tetrahydroxynaphthalene synthase, encoded by the *Streptomyces* rppA gene. This enzyme catalyzes the conversion of five malonyl-CoA molecules into one molecule of 1,3,6,8-tetrahydroxynaphthalene (THN) which is then spontaneously converted into the red-colored flaviolin in the presence of oxygen (Fig. 1A). The produced flaviolin gives a colorimetric indication of the intracellular malonyl-CoA levels that can be easily measured by means of absorbance. Furthermore, as the synthesis reaction of THN does not require any additional precursor such as ATP or NAD(P)H molecules, it is assured that this biosensor does not influence cell metabolism by altering energy or reducing power levels respectively. At first, the *Streptomyces coelicolor* A3(2) *rppA* was codon optimized based on *P. putida*’s GC-content and cloned under the control of the strong constitutive promoter BBa_J23100 in the medium copy vector pSEVAb23, resulting in plasmid pFLAV. After transformation into *P. putida*, the resulting LB plates containing *P. putida*-pFLAV appeared a dark-red color giving a first indication of successful expression of pFLAV (Fig. S1D). Having confirmed that *P. putida* expressing pFLAV produced flaviolin, we examined its potential as a dose-response biosensor. To characterize the dynamic response range of a biosensor, different concentrations of the target compound need to be added to the culture media. However, given that malonyl-CoA cannot be transported into the cell, we employed an indirect strategy using cerulelin. Cerulenin is a well-studied antibiotic isolated from various microorganisms which inhibits the fatty acid synthases systems. It has been shown in several organisms, including *Pseudomonas* species, that increasing cerulenin concentration gradually increases intracellular malonyl-CoA levels^1,2^. Thus, to evaluate the ability of the biosensor to respond to different malonyl-CoA concentrations, we cultured *P. putida*-pFLAV in vent cap 50 ml falcon tubes containing MOPS minimal medium supplemented with 70 mM glucose and various concentrations of cerulenin. As shown in Fig. 1B, the relative absorbance of flaviolin (OD_340_/OD_600_) from *P. putida*-pFLAV increased in line with the increasing concentration of cerulenin, whereas this did not apply to the control strain (*P. putida* WT carrying an empty pSEVAb23 vector). Therefore, we demonstrated the potential of pFLAV as a biosensor for high-throughput screening of *P. putida* malonyl-CoA overproducers.

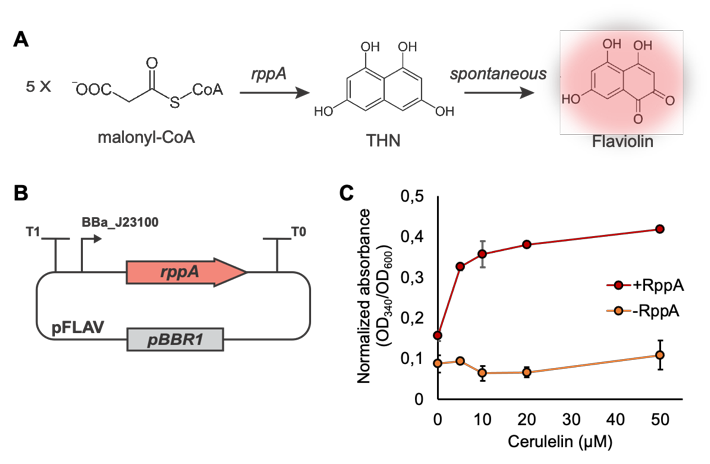

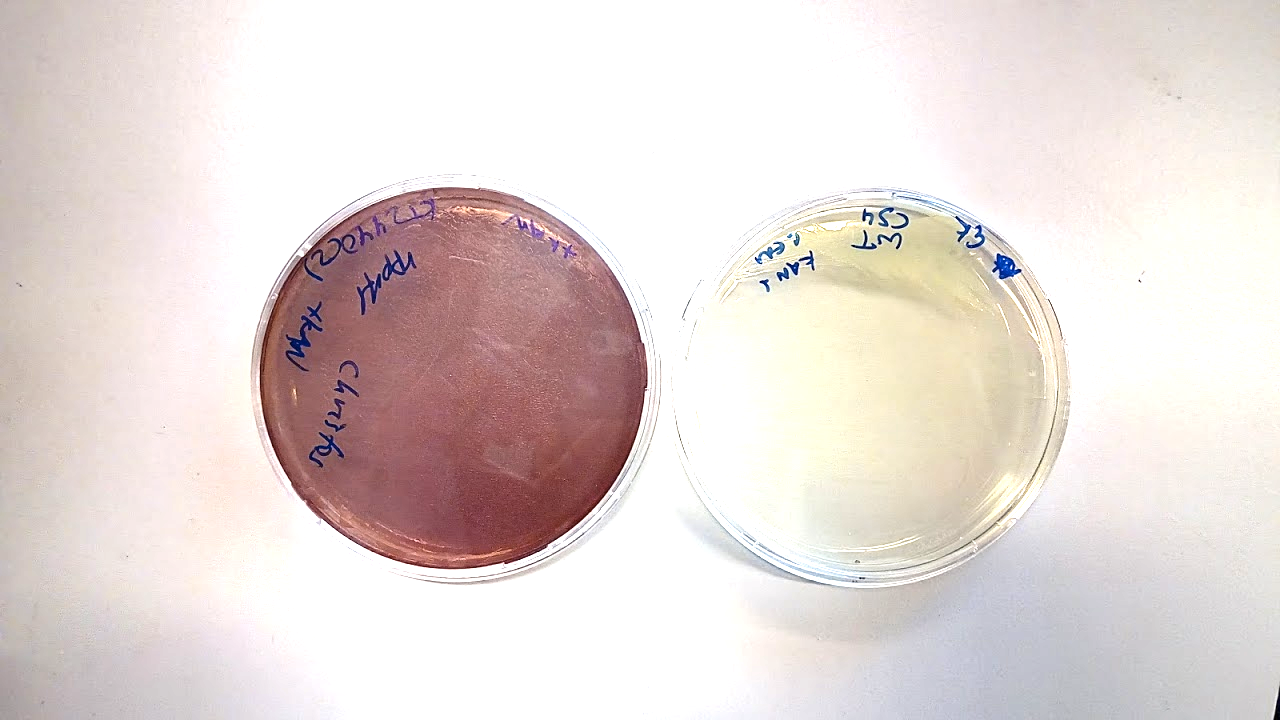

**D**

**Figure S3.** A) Synthesis of flaviolin from malonyl-CoA. First, five molecules of malonyl-CoA are converted into THN by RppA. Then THN is spontaneously converted to the red-colored flaviolin in the presence of oxygen. B) Graphic representation of the rppA expressing vector, pFLAV. rppA is under the control of BBa_J23100 promoter and flanked by the T1 and T0 terminator. C) Normalized flaviolin absorbance produce by the P. putida pFLAV grown on various concentration of cerulenin. D) P. putida transformed with pFLAV. Error bars indicate standard deviation of the data (n=3).

**Evaluation of the CRAGE-like Cre-mediated recombination system**

The utility of the presented CRAGE-like recombination system was initially demonstrated through the integration of two expression cassettes of different sizes. Two variants of pLGR were constructed carrying either a *sfGFP* gene or the four genes responsible for the production of protoviolacein (black pigment) *vioA*, *vioB*, *vioE* and *vioD*. Strains that have successfully incorporated the insert of interest into the target site should be gentamycin resistance as well as appear either fluorescent or black, thus allowing the simple evaluation of the recombination efficiency. After electroporation, the integration efficiency was calculated based on the colonies’ color to 100% and 90% for the *sfGFP* and the *vioABDE* operon, respectively. Indeed, colony PCR of the inserts revealed correct insert integration in all fluorescent and black colonies tested. Besides to integration efficiency, the total amount of the correctly engineered colonies is equally important since this method was intended to be used for the high-throughput integration of thousands of operon variants. Thus, a sufficient number of colonies per transformation is required for the proper operation of the tool. After electroporation the plates appeared full of colonies which could not be counted without further dilution. Even the larger *vioABDE* cluster was integrated at the same efficiency without affecting the transformation rate.

**REFERENCES**

1. Vance, D. *et al.* Inhibition of fatty acid synthetases by the antibiotic cerulenin. *Biochemical and Biophysical Research Communications* **48**, 649–656 (1972).
2. Kallscheuer, N., Vogt, M., Stenzel, A., Gätgens, J., Bott, M. & Marienhagen, J. Construction of a *Corynebacterium glutamicum* platform strain for the production of stilbenes and (2S)-flavanones. *Metabolic Engineering* **38,** (2016).
